# Allometric scaling of somatic mutation and epimutation rates in trees

**DOI:** 10.1101/2024.07.01.601331

**Authors:** Frank Johannes

## Abstract

How long-lived trees escape “mutational meltdown” despite centuries of continuous growth remains puzzling. Here we integrate recent studies to show that the yearly rate of somatic mutations and epimutations (μ_Y_) scales inversely with generation time (G), and follows the same allometric power law found in mammals (μ_Y_∝G^-1^). Deeper insights into the scaling function will permit predictions of somatic (epi)mutation rates from life-history traits without the need for genomic data.

---

Trees are among the longest living organisms on earth. With average lifespans of several hundred years and juvenile periods of up to half a century, they surpass the ontogenetic time-scales of most animal species by orders of magnitudes. Despite their extraordinary longevity, trees undergo continuous organogenesis throughout their lives, repeatedly forming lateral branches that give rise to leaves and fruits. The cell lineages that initiate branches derive from as few as 1-4 precursor cells of the shoot apical meristem (SAM)^1^. Repeated “sampling” of precursors during successive branching creates strong cellular bottlenecks and somatic drift^2,3^. This causes *de novo* mutations and epimutations that originate in the SAM to become fixed rapidly within clonal sectors of the tree topology^1^, and can thus be detected using sequencing approaches.

Recent genomic studies in trees found that the *per-year* rate of such fixed mutations and epimutations is surprisingly low (**Fig. 1A**). This observation has led to the hypothesis that long-lived trees evolved mechanisms to avoid the accumulation of potentially deleterious somatic variants with age. These types of protections are perhaps expected in long-lived plants, where - unlike in mammals - reproductive cells are formed from somatic precursors late in development^4^. Yet, substantial variation in somatic (epi)mutation rates exist between different tree species (**Fig. 1A**). To begin to understand the sources of this variation we analyzed somatic mutation and epimutation rates reported in 14 diverse tree species^5–15^ (**Table S1**). These include angiosperms (*Quercus robur, Prunus mira, Prunus persica, Prunus mume, Salix suchowensis, Populus trichocarpa, Shorea laevis, Shorea leprosula, Eucalyptus melliodora, Osmanthus fragrans, Citrus hybrid, Prunus hybrid, Fagus sylvatica*), as well as one gymnosperm (*Picea sitchensis*), and feature major differences in evolutionary histories (domesticated vs. natural), climate of origin (temperate vs. tropical climate), and life history traits (e.g. age at sexual maturity, and maximum lifespan).

**Figure 1:**
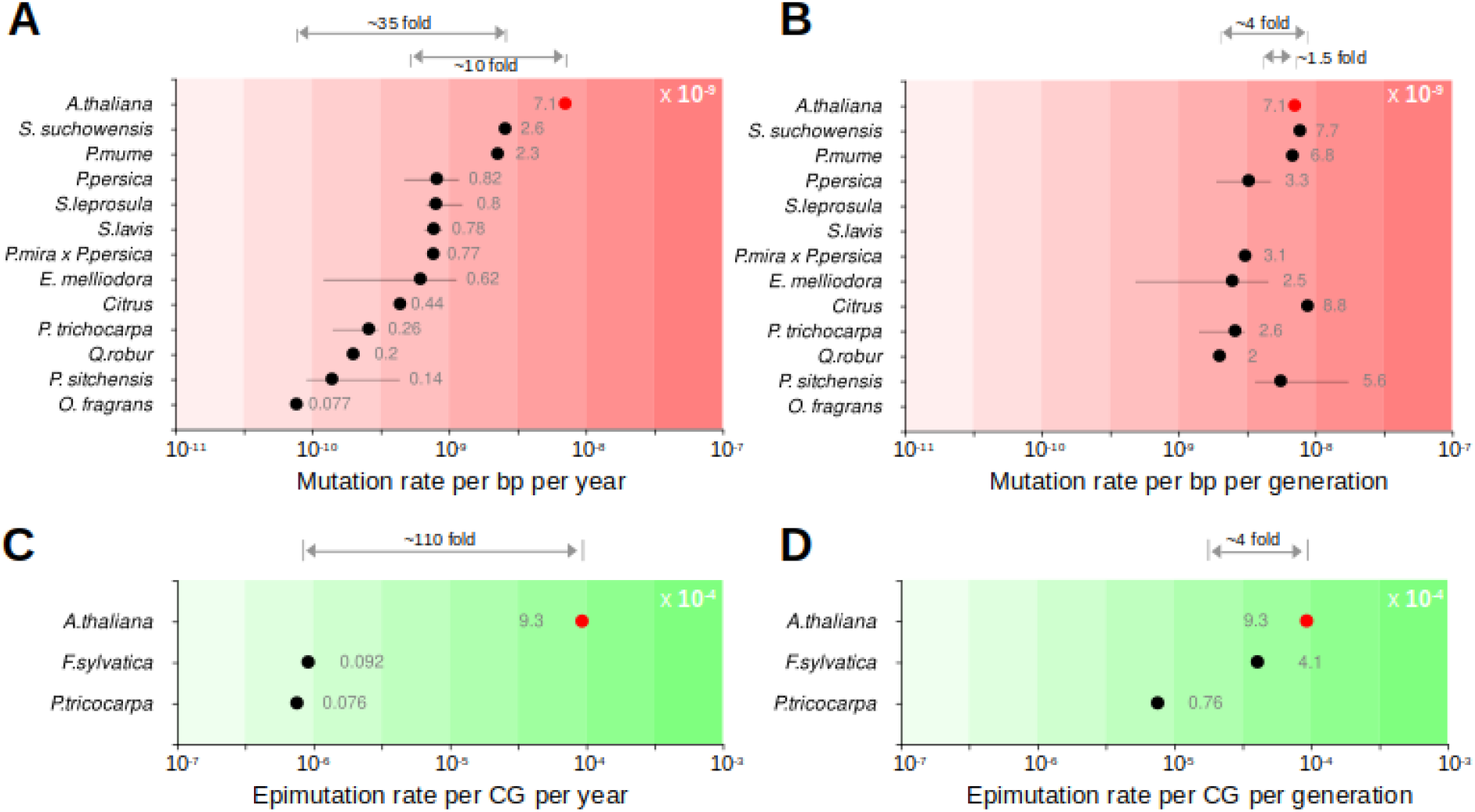
Variation in somatic mutation and epimutation rates across trees species. All rates were expressed as per diploid genome per bp and confidence intervals as +/- 1 SE of the estimate. **A**. Per-year somatic mutation rates for 12 tree species and the annual plant A. thaliana for comparison. **B**. Per generation mutation rates were obtained by multiplying the per year rates with generation time. **C**. Per-year somatic epimutation rates for 2 tree species and the annual plant A. thaliana. **D**. Per generation mutation rates were obtained by multiplying the per year rates with generation time.

A comparison of the *per-year* somatic mutation rates across species revealed that they vary approximately 35-fold, ranging from as low as 0.077 × 10^−9^ per diploid genome per bp (*O. fragrans*) to as high as 2.58 × 10^−9^ (*S. suchowensis*) (**Fig. 1A, Table S2**). On average, these rates were ∼10-fold lower than the estimated *per-year* mutation rate in the model plant *A. thaliana*^*16*^ (**Fig. 1A**). This shows that perennials incur fewer somatic mutations per unit time than annuals. Aging theory predicts that mutation rate variation is closely linked with life-history traits^17^. We found strong support for this theory. Indeed, despite major differences in experimental design and somatic variant calling approaches, inter-specific variation in *per-year* somatic mutation rates was inversely correlated with generation time in trees, and followed the same allometric power law linking somatic mutation rates and lifespan in mammals^18^ (**Fig. 2A-B, Methods**). This relationship has the simple form: μ_Y_ ≈ α·G^-1^, where μ_Y_ is the *per-year* (epi)mutation rate, G is generation time and α is a scaling constant. Remarkably, about 81% of the *per-year* rate variation among trees could be explained by differences in generation time, which compares to ∼82% being explained by lifespan in mammals^18^ (**Methods)**. This close resemblance illustrates that the relationship between mutation rates and life-history traits is highly conserved.

**Figure 2:**
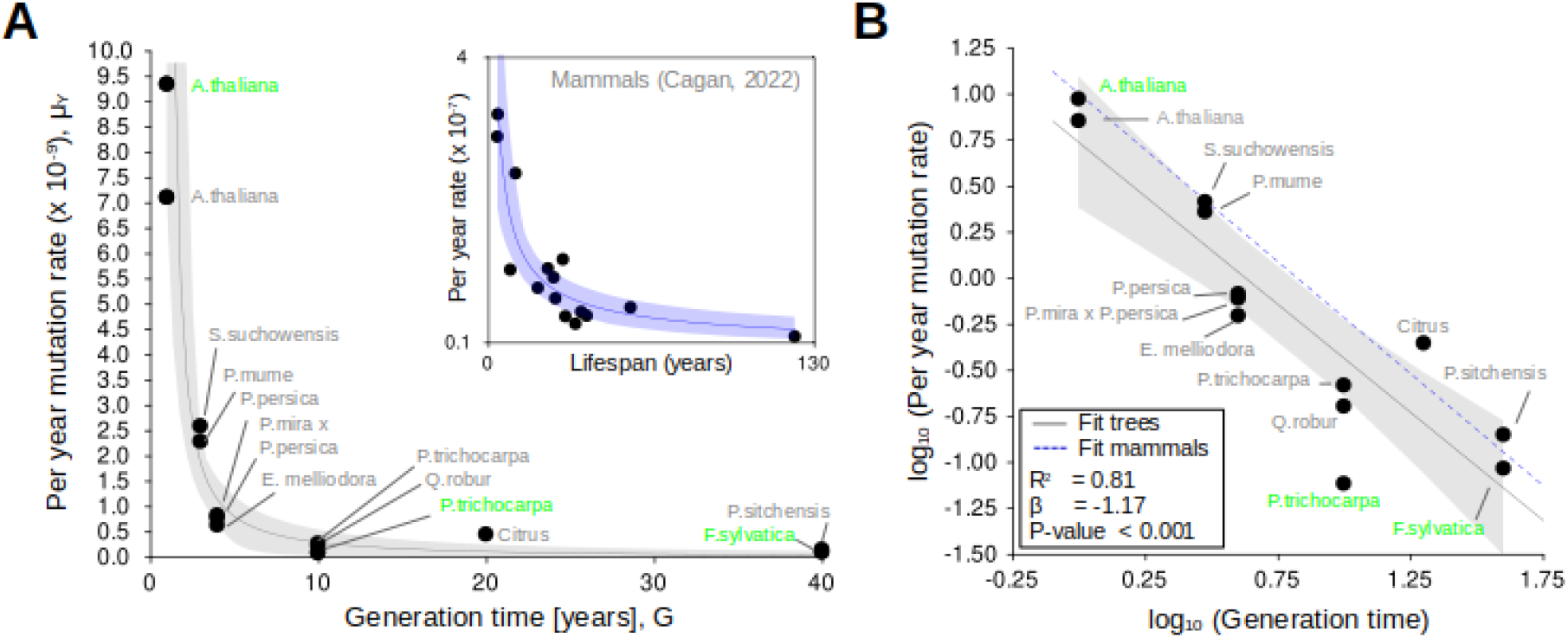
Somatic (epi)mutation rates scale with generation time. **A**. Generation time (G) is plotted against per year somatic mutation rates (µ). The power law function: µ = α ·G^β^ is fitted to the data (gray solid line) +/- SE (gray area). Inset: Re-analysis of mammalian somatic mutation rates as a function of lifespan. Data points labeled with green refer to scale- adjusted epimutation rates (i.e. epimuation rates multiplied by the constant 10^5^). **B**. The data in (A) plotted on a log_10_-log_10_ scale and fitted using linear regression. The re-scaled slope from the mammalian fit (blue dashed line) is superimposed.

The above insight becomes even clearer when re-scaling the *per-year* somatic mutation rates to *per-generation* rates. As a proxy, this can be done by simply multiplying the estimated *per-year* rates by the generation-time of each species^5^ (**Methods**). Doing this reduced rate variation among species from 35-fold to merely 4-fold, and placed the average mutation rate of trees (4.1 × 10^−9^) within close proximity of the *per-generation* rate of *A. thaliana* (7.1 × 10^−9^) (**Fig. 1B, Methods**). Similar observations could be made at the epigenetic level (**Fig. 1C-D**). Indeed, we found that estimates of the *per-year* somatic epimutation rates at CG dinucleotides in poplar and beech were 110-fold lower than the *per-year* rate in *A. thaliana*, but within 4-fold when adjusting for generation-time differences (2.4 × 10^−4^ per diploid genome per CG site vs. 9.3 × 10^−4^, respectively). Hence, the net generational burden of (epi)mutations is highly constrained across species.

Interestingly, the fact that the same scaling holds for both mutations and epimutations implies that their accumulation dynamics are modulated by the same underlying biology. In mammals, it has been argued that long-lived species have evolved more efficient DNA repair compared with short-lived species^18^. In trees with long generation times, improved DNA repair could counteract prolonged exposure to external mutagens, such as UV light. However, the DNA repair argument does not readily extend to CG epimutations in plants, as they are generally 4 orders of magnitude more frequent than DNA mutations^19^ (hence no 1:1 correspondence), have no reported link to DNA damage, and appear to be mainly due to methyltransferase errors during cell divisions^20^.

An alternative explanation is that long-lived tree species simply limit the number of meristematic cell divisions per unit time^1,4^. Although this hypothesis remains difficult to test in vivo, experimentally-induced growth acceleration in beech leads to an increase in xylem vessel, fiber and parenchyma cell divisions as well as a proportional increase in the rate of CG epimutations per unit time^14^. These latter observations are difficult to explain on the basis of differential DNA repair. The cell division hypothesis is consistent with the fact that lifespan and age at maturity are negatively correlated with growth rate^21^. It is therefore of theoretical interest to explore how the scaling 1/G (i.e. β=-1) connects with the fundamental allometric laws linking cellular metabolic rate, body size and growth discovered previously^22,23^. It would seem that this link only requires an auxiliary model that translates growth rates into meristematic cell divisions and fixed (epi)mutations per unit time.

Beyond theoretical considerations, the conserved allometry underlying (epi)mutation rate variation in trees also opens interesting practical possibilities. With precise estimates of the scaling function, it may be possible to accurately predict somatic (epi)mutation rates from knowledge of generation-times without the need for (epi)genomic measurements.

## Methods

### Processing of rate estimates

A total of 12 studies have reported somatic mutation and/or epimutation rates based on the analysis of 49 non-redundant trees, which correspond to 14 different tree species (**Table S1**). To make the reported rates more comparable, all rates were re-expressed as *per diploid genome per bp*. When necessary, reported 95% confidence intervals (95% CI) were converted to standard errors (SE) using: SE = 95% CI · 1.96^-1^. When multiple estimates were available for a given tree species, the average rate as well as the average SE was calculated. The large study by Wang et al.^6^ analyzed many replicate trees of the same species, but did not provide confidence intervals or standard errors. In this case, standard errors were calculated from the sample standard deviation (SD) using: SE = SD / *N*^-½^, where *N* is the number of replicate trees of a given species. Finally, only estimates based on leaf samples were taken forward. This only excluded three trees measured by Wang et al.^6^, which had data on root and flower pedals.

### Re-scaling per-year to per-generation (epi)mutation rates

A proxy for the *per generation* (epi)mutation rate (μ_*G*_) can be obtained by multiplying the estimated *per year* rate (μ_*Y*_) with the generation time (G) of a given species^5,8^:

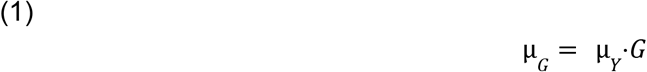

This calculation requires information about ‘generation time’. Formal population biology definitions of generation time combined knowledge of the growth rate of the population, survival probability (probability that an individual survives to a specific age), and age-specific fertility^24,25^. Many of these parameters are not easily obtainable in tree species, thus introducing substantial uncertainty when determining generation time. A more convenient measure, often also employed in the analysis of fruit trees, is ‘seed-to-first-seed’^26^, also known as ‘maturation age’^27^. Although this measure represents a lower-bound proxy of ‘generation time’ it does have the advantage that it is easy to ascertain and more readily extendable to tree species whose population ecology and life-histories are less well characterized. Information about see-to-first-seed generation time was available for 11 of 14 species from the references listed in **Table S2**.

### Allometric modeling and regression analysis

The nonlinear relationship between generation time (G) and the *per year* somatic mutation rate (μ_Y_) was modeled using the following power law function:

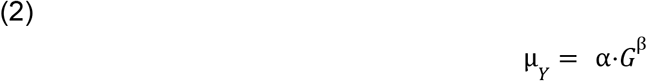

where β is the scaling exponent of the law. The basic structure of this law has been applied to a wide range of biological phenomena, for instance to model the relationships between body size and metabolic rate, or growth^22,28^, including in plants^29^. For **Fig. 2A**, parameter estimates and fits were obtained with the *nls()* package in R^30^. On a log-log scale, the above function is linear:

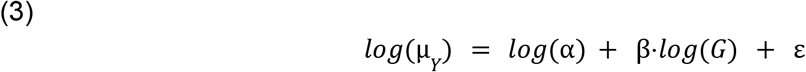

where we introduced ε∼ N(0, σ^2^), a normally distributed error term to account for measurement uncertainty. With the log-log transformed data, parameter estimation was performed with standard linear regression using the *lm()* function. The *R*^*2*^ statistic was used to quantify the proportion of variance in μ_Y_ which can be explained by the variance in G. An important property of the power law function in (2) is that it is scale invariant^22,28^. That is, if *h* and *k* are scaling constants along μ_Y_ and G, then

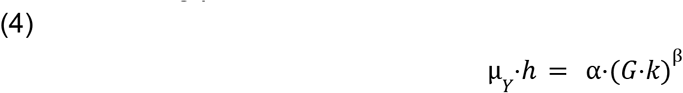

and

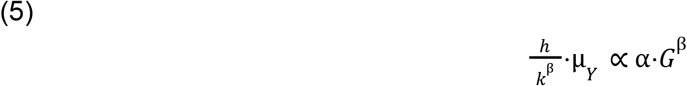

which means that inferences about the scaling exponent β are independent of the actual scale on which μ_Y_ and G are measured. In their allometric regression relating *per year* somatic mutation rates to lifespan in mammals, Cagan et al.^18^ employed the power law function in (2). They reported that the rate parameter β is not significantly different from -1. Their finding suggests that the *per year* somatic mutation rate is inversely proportional to lifespan. To test if this allometric property also holds in trees we performed a one-sample T-test about the regression slope. The test statistic has the form:

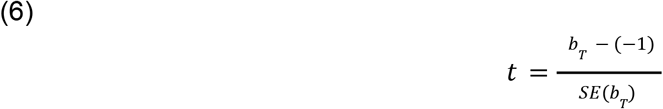

where *b*_*T*_ is the estimated regression slope. The degrees of freedom of the test are *N*_*T*_ - 2, with *N*_*T*_ being the sample size of the tree data. We tested H_0_: β = -1 against H_A_: β ≠ -1, and were unable to reject the null hypothesis (*t*_*11*_ = 0.99, *p* = 0.17). Hence, consistent with mammals, the *per year* somatic (epi)mutation rate is inversely proportional to ‘generation time’

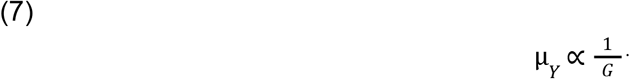

This inverse scaling thus appears to be a conserved property of (epi)mutation rate evolution.

### Co-visualization of the mammalian and tree data

Measurements of *G* and μ_Y_ exist on a different scale in mammals compared to trees. For **Fig. 2B**, we sought to visualize the regression slopes in the same plot. To do this, we re-analyzed the data from Cagan et al.^18^ using the same analysis procedure used for the tree data. We calculated the *per year* somatic mutation rate for each mammalian species using μ_Y_ = *S*·*L*^*-1*^, where *S* is the mean number of substitutions per year (termed ‘Mean mutation rate (SBS/genome/year)’ in their data) and *L* is the total number of callable sites in the genome (termed ‘Analyzable genome size (bp)’ in their data). We then normalized G and μ_Y_ using:

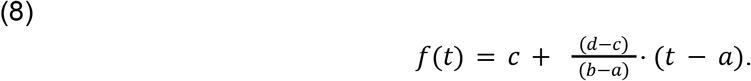

This function maps a value *f* from the interval [*a, b*] of mammalian data into the interval [*c, d*] of the tree data while preserving the relative ranking of the data points. Here a, c and b, d are the minimum and maximum of the respective data ranges. The normalization was carried out for the *G* and μ_Y_ variables separately. Finally, we estimated the scale parameter β based on this normalized mammalian data, which we visualized in **Fig. 2B** (blue dashed line).

## Acknowledgements

I thank H. Pretzsch, T. Reusch and H. Flachowsky for feedback, as well as G. Schmied, T. Hilmers and H. Flachowsky for their help in finding resources. This work was supported by the Bundesministerium für Bildung und Forschung (Project: epiSOMA).

