## Supplementary material for "Allometric scaling of somatic mutation and epimutation rates in trees": Table S1

**Table S1: Overview of studies and tree species reporting somatic mutation and epimutation rates in trees.**

| Focus | Year | Author | Species | Clade | Taxa | Genome (Mb) | Tree | Age of sampled tree | Target tissue | N samples |
|---|---|---|---|---|---|---|---|---|---|---|
| Mutation | 2017 | Schmid-Siegert et al. | Oak (Quercus robur) | Angiosperm | Fagaceae | 740 | 1 | 234 | Leaf | 26 |
| Mutation | 2019 | Wang et al. | Peach (Prunus mira) | Angiosperm | Rosaceae | 225 | 1 | 600 | Leaf | 32 |
| Mutation | 2019 | Wang et al. | Peach (Prunus mira) | Angiosperm | Rosaceae | 225 | 2 | 550 | Leaf | 12 |
| Mutation | 2019 | Wang et al. | Peach (Prunus mira) | Angiosperm | Rosaceae | 225 | 3 | 420 | Leaf | 23 |
| Mutation | 2019 | Wang et al. | Peach (Prunus mira) | Angiosperm | Rosaceae | 225 | 4 | 300 | Leaf | 9 |
| Mutation | 2019 | Wang et al. | Peach (Prunus persica) | Angiosperm | Rosaceae | 225 | 1 | 21 | Leaf | 23 |
| Mutation | 2019 | Wang et al. | Peach (Prunus persica) | Angiosperm | Rosaceae | 225 | 1 | 21 | Root | 13 |
| Mutation | 2019 | Wang et al. | Peach (Prunus persica) | Angiosperm | Rosaceae | 225 | 2 | 25 | Leaf | 16 |
| Mutation | 2019 | Wang et al. | Peach (Prunus persica) | Angiosperm | Rosaceae | 225 | 2 | 25 | Pedal | 13 |
| Mutation | 2019 | Wang et al. | Peach (Prunus persica) | Angiosperm | Rosaceae | 225 | 3 | 30 | Leaf | 26 |
| Mutation | 2019 | Wang et al. | Peach (Prunus persica) | Angiosperm | Rosaceae | 225 | 4 | 50 | Leaf | 6 |
| Mutation | 2019 | Wang et al. | Peach (Prunus persica) | Angiosperm | Rosaceae | 225 | 5 | 40 | Leaf | 16 |
| Mutation | 2019 | Wang et al. | Peach (Prunus persica) | Angiosperm | Rosaceae | 225 | 6 | 2 | Leaf | 75 |
| Mutation | 2019 | Wang et al. | Plum (Prunus mume) | Angiosperm | Rosaceae | 220 | 1 | 20 | Leaf | 25 |
| Mutation | 2019 | Wang et al. | Plum (Prunus mume) | Angiosperm | Rosaceae | 220 | 1 | 20 | Root | 32 |
| Mutation | 2019 | Wang et al. | Plum (Prunus mume) | Angiosperm | Rosaceae | 220 | 2 | 8 | Leaf | 33 |
| Mutation | 2019 | Wang et al. | Shrub willow (Salix suchowensis) | Angiosperm | Salicaceae | 480 | 1 | 1 | Leaf | 19 |
| Mutation | 2019 | Wang et al. | Shrub willow (Salix suchowensis) | Angiosperm | Salicaceae | 480 | 1 | 1 | Root | 21 |
| Mutation | 2020 | Hofmeister et al. | Poplar (Populus trichocarpa) | Angiosperm | Salicaceae | 390 | 1 | 330 | Leaf | 8 |
| Mutation | 2023 | Satake et al. | Balau (Shorea laevis) | Angiosperm | Dipterocarpaceae | 347 | 1 | 324 | Leaf | 7 |
| Mutation | 2023 | Satake et al. | Balau (Shorea laevis) | Angiosperm | Dipterocarpaceae | 347 | 2 | 187 | Leaf | 7 |
| Mutation | 2023 | Satake et al. | Red Meranti (Shorea leprosula) | Angiosperm | Dipterocarpaceae | 376 | 1 | 78 | Leaf | 7 |
| Mutation | 2023 | Satake et al. | Red Meranti (Shorea leprosula) | Angiosperm | Dipterocarpaceae | 376 | 2 | 54 | Leaf | 7 |
| Mutation | 2020 | Orr et al. | Yellow box (Eucalyptus melliodora) | Angiosperm | Myrtaceae | 550 | 1 | guess: 50-200 | Leaf | 24 |
| Mutation | 2019 | Hanlon et al. | Sitka spruce (Picea sitchensis) | Gymnosperm | Pinaceae | 21000 | 20 | range: 220-500 | Leaf | 80 |
| Mutation | 2022 | Duan et al. | Sweet olive (Osmanthus fragrans) | Angiosperm | Oleaceae | 780 | 1 | 1700 | Leaf | 10 |
| Mutation | 2022 | Perez-Roman et al. | Citrus (Citrus x clementina hort. ex Tanaka) | Angiosperm | Rutaceae | 300 | 1 | 36 | Leaf | 15 |
| Mutation | 2016 | Xie et al. | Peach (Prunus mira x persica) | Angiosperm | Rosaceae | 225 | 1 | parent-offspring | Leaf | 70 |
| Mutation | 2011 | Ossowski et al. | A. thaliana | Angiosperm | Brassicaceae | 125 | | 1 | Leaf | |
| Epimutation | 2024 | Zhou et al. | European beech (Fagus sylvatica) | Angiosperm | Fagaceae | 550 | 1 | 200 | Leaf | 10 |
| Epimutation | 2024 | Zhou et al. | European beech (Fagus sylvatica) | Angiosperm | Fagaceae | 550 | 2 | 200 | Leaf | 7 |
| Epimutation | 2020 | Hofmeister et al. / Shahryary et al. | Poplar (Populus trichocarpa) | Angiosperm | Salicaceae | 390 | 1 | 330 | Leaf | 8 |
| Epimutation | 2023 | Yao et al. | A. thaliana | Angiosperm | Brassicaceae | 125 | | 1 | Leaf | |
