## Supplementary material for "Allometric scaling of somatic mutation and epimutation rates in trees": Table S2

**Table S2: Estimates of somatic mutation and epimutation rates.**

| Focus | Species | Rate estimate | SE (lower) | SE (upper) | Generation time |
|---|---|---|---|---|---|
| Mutation | O. fragrans | 0.077 | 0.069 | 0.085 | |
| Mutation | P. sitchensis | 0.14 | 0.09 | 0.43 | 40 |
| Mutation | Q.robur | 0.2 | 0.18 | 0.22 | 10 |
| Mutation | P. trichocarpa | 0.26 | 0.14 | 0.3 | 10 |
| Mutation | Citrus | 0.44 | 0.44 | 0.44 | 20 |
| Mutation | E. melliodora | 0.62 | 0.12 | 1.12 | 4 |
| Mutation | P.mira x P.persica | 0.77 | 0.69 | 0.81 | 4 |
| Mutation | S.lavis | 0.775 | 0.655 | 0.88 | |
| Mutation | S.leprosula | 0.805 | 0.68 | 1.235 | |
| Mutation | P.persica | 0.817 | 0.468 | 1.165 | 4 |
| Mutation | P.mume | 2.275 | 2.17 | 2.38 | 3 |
| Mutation | S. suchowensis | 2.58 | 2.58 | 2.58 | 3 |
| Mutation | A.thaliana | 7.1 | 6.4 | 7.8 | 1 |
| Epimutation | P.tricocarpa | 0.076 | 0.076 | 0.076 | 10 |
| Epimutation | F.sylvatica | 0.092 | 0.082 | 0.102 | 40 |
| Epimutation | A.thaliana | 9.337 | 7.071 | 11.603 | 1 |

**Note:** Mutation rates are given as x 10^-9 and epimutation rates as x 10^-4
